# Activity Protects Spinal Premotor Interneurons from Microglial Phagocytosis and Transneuronal Degeneration After Corticospinal Injury

**DOI:** 10.1101/2025.02.24.639981

**Authors:** Jasmine Pathan, Alzahraa Amer, Yutaka Yoshida, John H. Martin

**Affiliations:** Department of Molecular, Cellular, and Biomedical Sciences, Center for Discovery and Innovation, City University of New York School of Medicine, New York, NY 10032, USA; Neural Connectivity Development in Physiology and Disease Laboratory, Burke Neurological Institute, White Plains, NY 10605, USA; Brain and Mind Research Institute, Weill Cornell Medicine, New York 10065, USA; Neural Circuit Unit, Okinawa Institute of Science and Technology Graduate University, Onna-son, Kunigami-gun, Okinawa 904-0495, Japan; Neuroscience Program, Graduate Center of the City University of New York, New York, NY 10016, USA

## Abstract

Spinal premotor circuits play a fundamental role in motor control. The corticospinal tract (CST) provides control signals to premotor circuits in the spinal cord, guiding voluntary skilled movements. Unilateral selective lesion of the CST in the medullary pyramidal tract (PTX) produces transneuronal degeneration, whereby Choline Acetyltransferase-positive (ChAT) premotor interneurons contralesionally undergo non-apoptotic degeneration by microglial phagocytosis. Evidence shows that transneuronal degeneration has an activity dependence: MCX inactivation produces transneuronal degeneration and spinal DC neuromodulation after PTX ameliorates it. This study expands our understanding of transneuronal degeneration mechanisms by examining the activity-dependence of degeneration vulnerability and the implications for motor recovery in a mouse model of a complete CST lesion model (bilateral PTX). We address four key unanswered questions: Are Chx10 (VGlut2) interneurons, the largest spinal interneuron class to receive direct synaptic connections from the CST vulnerable to transneuronal degeneration after CST loss; using DREADD neuromodulation, what are effective sources of presynaptic activation for protecting spinal premotor interneurons after CST loss; how are ameliorating transneuronal interneuron degeneration and reducing microglial activation associated; and does effective rescue of interneuron degeneration rescue grip strength after injury? Transneuronal degeneration is a pervasive pathophysiological change injury; CST lesion produces significant Chx10 interneuron loss. Multiple sources of neuronal DREADD activation—motor cortex, reticular formation, and spinal interneurons—are effective in ameliorating transneuronal degeneration. Interneuron rescue is strongly associated with ameliorating inflammation, showing potential causality between interneuron degeneration and inflammation after CST lesion. Finally, rescuing spinal interneurons was associated with restoring function. Our findings demonstrate the interplay between neuronal activity, microglia actions mediating transneuronal degeneration, and motor recovery following CNS injury.

## Introduction

Spinal premotor circuits play a fundamental role in motor control by integrating inputs from supraspinal centers in the cortex and brain stem with peripheral sensory feedback, to regulate motor neuron activity and coordinate muscle contractions (Bareyre et al., 2005; Ueno et al., 2018). The corticospinal tract (CST), which originates predominantly from the motor cortex (MCX), provides key control signals to premotor circuits in the spinal cord, guiding voluntary skilled movements (Lemon, 2008).

Lesion of the CST produces transneuronal degeneration, whereby spinal interneurons undergo degeneration after loss of this important presynaptic input. Selective unilateral CST lesion of the pyramidal tract (PTX) produces transneuronal degeneration of Choline Acetyltransferase-positive (ChAT) premotor interneurons contralesionally. Mediated through non-apoptotic microglial phagocytosis (phagoptosis; (Brown and Neher, 2014; Brown et al., 2015)), there is both a rapid loss of their cell bodies and the loss of their motor neuron synapses (C-boutons)(Jiang et al., 2018). Transneuronal degeneration, observed in our lesion-based study, has an activity dependence: MCX pharmacological inactivation also produces transneuronal degeneration, while increasing neural activity through spinal DC neuromodulation after PTX prevents it.

This study expands our understanding of transneuronal degeneration mechanisms by examining the activity-dependence of interneuron degeneration vulnerability and the implications for motor recovery. We used a complete CST lesion model (bilateral pyramidal tract section, biPTX) in the mouse. A unilateral CST lesion spares ipsilateral projections from the contralateral hemisphere (Brosamle and Schwab, 2000), whose presence may rescue CST interneuron targets after injury. Using this complete bilateral CST lesion model, we addressed four key unanswered questions. **First**, are other interneuron classes vulnerable to degeneration after CST loss? We hypothesized that Chx10 (VGlut2) interneurons, the largest spinal interneuron class to receive direct synaptic connections from the CST (Ueno et al., 2018), are vulnerable to transneuronal degeneration after CST loss. **Second**, are other sources of presynaptic drive to spinal premotor interneurons, in addition to the CST, capable of supporting interneuron viability after CST loss? We used rigorous DREADD-based approaches (Roth, 2016) to target neuronal activity modulation to specific cell populations in the MCX, reticular formation, and the cervical enlargement, to determine if neural activation by these sources could rescue ChAT and Chx10 interneurons after a complete CST lesion. **Third**, how do microglia respond to surrogate sources of activity in modulating transneuronal interneuron degeneration (Jiang et al., 2018)? **Finally**, as premotor interneurons are key to a wide-range of motor behaviors (Zagoraiou et al., 2009; Dougherty and Kiehn, 2010; Hagglund et al., 2013), what is the association between interneuron rescue by providing activity after injury, and grip strength recovery?

Our study reveals that transneuronal degeneration is a pervasive pathophysiological change injury, impacting major interneuron classes mediating CST premotor control. Multiple sources of neuronal DREADD activation—motor cortex, reticular formation, and spinal interneurons—were effective in ameliorating transneuronal degeneration. Rescue of interneurons was robustly associated with ameliorating inflammation, showing potential causality between interneuron degeneration and inflammation after CST lesion. Finally, rescuing spinal interneurons was associated with restoring function. Our findings demonstrate the interplay between neuronal activity, microglia actions in transneuronal degeneration, and motor function recovery following CNS injury-induced neural activity loss.

## Materials and Methods

### Animals

Male and female C57BL/6J, Pitx2-Cre, Chx10-Cre, and VGlut2-Cre adult mice (4-7 months old) were on a C57BL/6J background. Mice were group-housed on a 12-hour light/dark schedule with ad libitum access to food and water. All animal protocols and procedures were approved by the Institutional Animal Care and Use Committee (IACUC) at the City College of New York. All studies are in accordance with the National Institutes of Health Guide for the Care and Use of Laboratory Animals.

All mice were randomly placed into the following six groups. (1) The naïve group received no treatment and was housed as described above until perfusion. (2) The bilateral pyramidal tract lesion (biPTX) group received a complete CST lesion and was observed for ten days post-injury (dpi). (3) After establishing activity-dependence, we used the biPTX model but used excitatory motor cortex DREADDs (biMCX Gq) to investigate if neural activity could act through cortico-brain stem-spinal pathways to prevent transneuronal degeneration. (4) To further confirm this hypothesis, we used a VGlut2 Cre mouse line for cell-specific activation of the medullary reticular formation, including MdV after a biPTX (biMdV Gq). (5) To test whether the activity could bypass the CST completely and utilize local spinal circuits to prevent transneuronal degeneration, we used a biPTX model with excitatory spinal DREADDs (biSC Gq). The biMCX Gq, biSC Gq, and biMdV Gq groups received biPTX at the appropriate timepoint of transfection (MCX 3 weeks; MdV 2 weeks; SC 2 weeks) and were euthanized 10dpi. (6) A biPTX group that did not receive DREADD treatment was given ten days of clozapine nitric-oxide (CNO) to confirm that CNO did not have off-target effects and influence grip strength.

### Surgical and Stereotaxic Injection Procedures

Mice were anesthetized with 5% isoflurane and maintained under anesthesia with 3% isoflurane for the duration of the surgery. The depth of anesthesia was confirmed via the absence of a toe pinch test. Body temperature was maintained at 37°C with a heating pad. Eye lubricant was applied to both eyes (Paralube Vet Ointment). The mouse was fixed in a stereotaxic frame with ear bars and a mouthpiece adaptor (David Kopf Instruments). Pulse oximetry was monitored throughout the surgical procedure. Following the surgical procedure, the skin was sutured, and an analgesic was administered (Buprenorphine (SQ); 0.1mg/kg). Animals were then placed in a recovery cage on a heating pad and observed until ambulatory.

### Bilateral Pyramidotomy

The animal was placed in a supine position, and a midline skin incision was made to the neck. The underlying muscles were blunt dissected until the trachea was exposed. The trachea was displaced to expose the ventral occipital bone. A craniotomy was made using rongeurs to expose the medullary pyramids(Serradj et al., 2023). The dura was cut using iridectomy scissors. Using a microscalpel, the medullary pyramids were bilaterally lesioned ∼0.5mm deep while ensuring no damage to the basilar artery. Absorbable gelatin powder and saline were used to clean the injury site prior to superficial skin suturing.

### Motor Cortex DREADD AAV Injections

DREADDs were used to modulate neuronal activity (Roth, 2016; Amer and Martin, 2022). A midline incision and a bilateral craniotomy were made over the skull, exposing the forelimb motor cortex of C57BL/6J mice. A total of six DREADD viral injections (AAV2-CamKII-hM3Dq-mCherry or AAV2-CamKII-hM4Di-mCherry) were performed using a stereotaxic injection system. The virus was injected using a 35-gauge needle at a rate of 300nl min^-1^, and the needle was held in place for two minutes post-injection to prevent backflow. All motor cortex injections were based on stereotaxic coordinates that evoke a motor response (Asante and Martin, 2013). Injection sites, relative to bregma, were made at 1) 1.0 mm anteroposterior (AP), ± 0.9 mm mediolateral (ML), 2) 1.5 mm anteroposterior (AP), ± 1.0 mm mediolateral (ML), 3) 1.2 mm anteroposterior (AP), ± 1.9 mm mediolateral (ML), and 0.9mm dorsoventral (DV). Following the injections, the superficial skin was sutured.

### Spinal Cord Injections

The following mouse lines were used in this procedure: C57BL/6J, Pitx2-Cre, and Chx10-Cre. A midline incision was made from T2 to the base of the skull, and the dorsal neck muscles were bluntly dissected to expose the cervical vertebrae from C2 to T2. The spiny process of T1 was clamped with an alligator clip held by a spinal vertebrae clamp (David Kopf Instruments), which was used to gently elevate the spinal cord. Laminectomies were performed at the C5 and C7 segments. The dura was removed with forceps, and the animal received three bilateral spinal injections (a total of six injections) per segment. DREADD virus (AAV2-CamKII-hM3Dq-mCherry for the wild-type mice; AAV2-hSyn-DIO-hM3Dq-mCherry for the Cre-lines) was injected using a glass micropipette (40um diameter) using the same parameters as described above. Injection sites were made 1) ± 0.5 mm mediolateral (ML) and 2) 0.8 mm dorsoventral (DV) to target the intermediate zone at the level of the central canal(Serradj et al., 2014). For anteroposterior (AP) injections, the most rostral and caudal part of the segments were determined for the first two injections, and the midpoint between those two sites was the third injection site. Following the injections, the skin was sutured in two layers: the deep tissue later with re-absorbable sutures and the superficial skin with non-re-absorbable sutures.

### Brain Stem Injections

An incision was made over the skull, exposing the cerebellum for a bilateral craniotomy to expose the injection sites in the VGlut2 Cre mouse line. Two DREADD viral injections (AAV2-hSyn-DIO-hM3Dq-mCherry) were performed using a stereotaxic injection system.

DREADD virus was injected using the same parameters as the motor cortex injections. Injection sites, relative to bregma, were made at -5.8 mm anteroposterior (AP), ± 0.4 mm mediolateral (ML), and -5.2 mm dorsoventral (DV) (Esposito et al., 2014). Following the injections, the superficial skin was sutured.

### Clozapine N-Oxide Intraperitoneal Injections

To activate the DREADDs injected into the regions described above, intraperitoneal injections of clozapine N-oxide (CNO; 2 mg/kg; i.p.) were performed daily for ten consecutive days, 24 hours after CST injury. Grip strength was obtained during the washout period and one hour after CNO i.p. injections to confirm DREADD activation.

### Forelimb Grip Strength Test

The grip strength of each mouse (Anderson et al., 2004)was measured using a grip strength apparatus (Columbus Instruments). Mice would grip a horizontal bar attached to a force transducer, which measured the maximum force (peak tension force) the mouse applied during the pull. The force transducer was set to peak tension mode with grams as a unit of value. We utilized a published protocol (Aartsma-Rus and van Putten, 2014): The mouse was weighed to normalize for body weight before the test. The mouse was then held by the base of the tail and moved horizontally towards the bar. As they approach the bar, the mouse reaches and grabs the bar and pulls. After three pulls (one trial), the mouse returned to the cage for a one-minute resting period. A total of five trials were performed (fifteen pulls). The highest three pulls were averaged and divided by the mouse’s body weight. Mice were assessed before stereotaxic surgical procedures to obtain a baseline and verify that DREADD injections in the motor cortex, spinal cord, or brain stem did not affect behavior. Mice were assessed on the day of bilateral pyramidal tract lesions before the injury, on days 1, 3, and 5 after the injury, and on the day of perfusion. Systemic administration of CNO increases neuronal activity for approximately 1.5 hours, following which activity returns to baseline (Yang and Martin, 2023). We measured the peak tension force before CNO administration to prevent the activity boost during this period of CNO-mediated DREADD action from increasing muscle strength.

### Tissue Preparation and Immunohistochemistry

Animals were anesthetized with ketamine/xylazine injections (ketamine 100mg/kg; xylazine 10mg/kg, i.p.) and perfused intracardially with normal 0.9% saline followed by 4% PFA. The brain and spinal cord were removed, postfixed in 4% PFA for 30 minutes, then transferred to a 30% sucrose solution for 24 hours or until the tissue sunk. The frozen spinal cord was sectioned into 40um-thick coronal sections using a sliding microtome. Six sections from the cervical enlargement were selected from each animal. Free-floating sections were washed three times in 1XPhosphate-buffer saline (PBS) and then incubated in a blocking buffer (1XPBS with 0.2% Tween, 3% donkey serum) on a shaker at room temperature for 1 hour. Primary antibody (refer to Key Resources Table) was added to the blocking buffer and the tissue was incubated at 4°C overnight. Sections were washed three times in 1XPBS before secondary antibody (1:1000 Alexa Fluor 448 donkey anti-goat, 1:1000 Alexa Fluor 448 donkey anti-rat, 1:1000 Alexa Fluor 561 donkey anti-rabbit, 1:1000 Alexa Fluor 561 donkey anti-mouse) was added and incubated for 2 hours at room temperature. Sections were washed three times in 1xPBS before being mounted with Vectashield and stored in the dark at 4°C until imaging.

### Confocal Imaging

Confocal images were taken with an LSM 880 microscope (Carl Zeiss). Montage images and z-axis stacked images were taken with a motorized stage. The laser wavelengths were as follows: 488, 561, and 631. Bilateral images were captured as z-stacks with a 1.5um step size at 40x magnification with 354.25 um x 354.25 um dimensions within the medial intermediate zone, adjacent to the central canal, and ventral to the ventral border of the dorsal column. This region was chosen due to the known expression of Pitx-2, the transcription factor for a subset of ChAT interneurons(Zagoraiou et al., 2009). The z-stacks were opened in Fiji (NIH image analysis software, http://imagej.nih.gov/ij/), and each cell was marked using the ‘Cell Counter’ function. Montage images were taken at 20x magnification for heatmaps to encompass the entire spinal section. Images were taken from five to eight animals from each group and five sections from each animal.

### Spinal Interneuron and Microglia Quantification

Changes in ChAT and Chx10 premotor interneurons, microglia, C1q+ neurons, mCherry+ neurons, and cFos+ neurons were determined by the absolute counts of cells within the region of interest (ROI) through immunolabeling. ChAT and Chx10 interneurons were identified using colabeling of NeuN and ChAT or NeuN and Chx10. Phagocytic microglia were identified with co-immunolabeling of Iba-1, lysosomal marker CD68, and DAPI. C1q positive neurons were determined through colabeling of C1q and NeuN. The DREADD virus used was coupled with the fluorophore mCherry to identify DREADD transfected cells. We identified mCherry positive neurons endogenous mCherry expression (intranuclear and cytoplasmic labeling) from DREADD transfection. We identified and quantified co-labeled mCherry+NeuN+ positive immunostained neurons in addition to mCherry+ChAT+, and mCherry+Chx10+ premotor interneurons. We confirmed that excitatory DREADDs increased ChAT and Chx10 interneuron activity by immunostaining for cFos.

### Interneuron Density Heat Maps

ChAT and Chx10 premotor interneurons were counted as described above. A semi-automatic analysis was employed for heatmap generation and cell quantification using image analysis software Fiji. First, a region of interest was manually selected within the gray matter of spinal sections. Cells positive for ChAT or Chx10 were marked at the corresponding location of the cells using the “Cell Counter” function in Fiji. Images were generated into an 8-bit binary mask after thresholding to segregate the marked cells from the selected region of interest. For each sample, a heatmap was constructed to determine cell density and distribution of ChAT and Chx10 premotor interneurons using R Studio. Using the packages ggPlot2 and reshape2, heatmaps were generated. Each group consisted of 5-6 animals, and 6 sections per group were selected for heat mapping.

### Statistics

All animals were randomly placed into the groups described above. All cell counts were performed by lab personnel blinded to the animal group. An a priori power analysis was conducted with G*Power. Statistical analysis was performed using GraphPad Prism (GraphPad Software, Version 6.0, La Jolla, CA, United States). For all cell counts, there was a normal distribution; an ordinary one-way ANOVA with Bonferroni’s correction for multiple comparisons was performed. Data are presented as mean ± standard error of the mean with p < 0.05 considered as statistically significant.

## Results

### Experimental Design

We examined the effects of DREADDs in MCX, brain stem, or spinal cord on transneuronal degeneration of ChAT and Chx10 premotor spinal interneuron (IN) classes (Fig 1A). Using cFos immunohistochemistry as a validation method revealed a clear histological (Fig 1B) and statistically significant enhancement (Fig 1C-D) in DREADD-transduced neuron activity in layer five of the motor cortex following systemic administration of CNO. Using the cFos assay (31.4% increase) demonstrates the effectiveness of CNO injections in stimulating neuronal activity (Fig 1C-D, p=0.0350).

**Figure 1:**
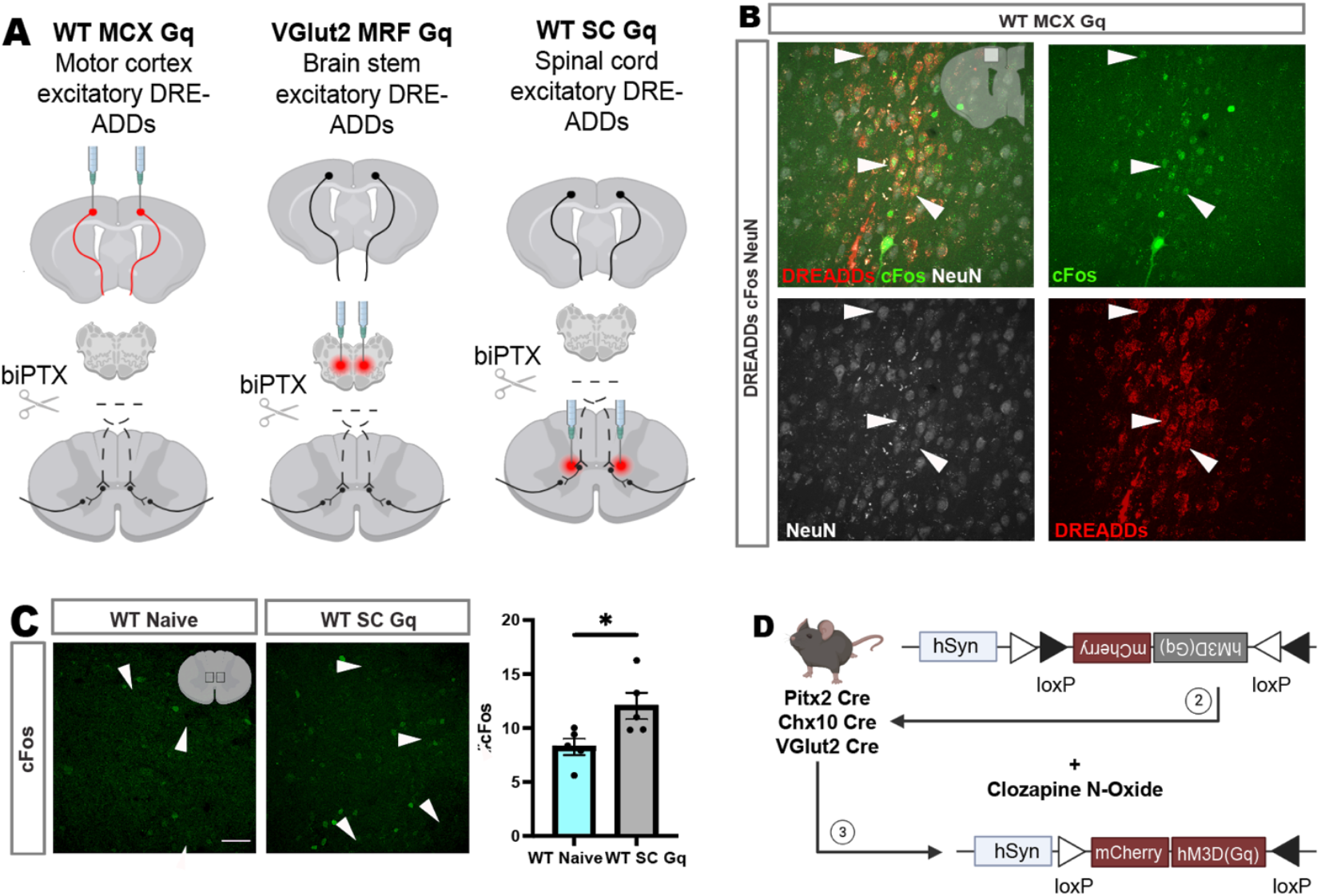
Experimental design to examine the effects of DREADDs on transneuronal degeneration of ChAT and Chx10 premotor spinal interneurons (IN). (A) Schematic representations show the use of excitatory DREADDs in these regions. Mice received bilateral injections of DREADD virus (AAV2-CamKIIa-hM3D(Gq)-mCherry) into the MCX forelimb region or the cervical segments of the spinal cord, or AAV2-hSyn-hM3D(Gq)-mCherry into specific brain stem or spinal cord regions in Cre mice. All mice were administered 10 daily injections of clozapine N-oxide (CNO) starting at least 14 days post-injection. (B) Immunohistochemical analysis using cFos as a marker of neuronal activity. Representative images demonstrate increased cFos expression in DREADD-transfected neurons, indicating enhanced neuronal activity after CNO administration. (C) Quantification of cFos-positive cells shows a statistically significant increase in neuronal activity in DREADD-transduced neurons compared to wild-type (WT) naïve controls (p=0.0350). Error bars represent the standard error of the mean (SEM), and each data point represents biological replicates with 10 technical replicates per sample. (D) Schematic representation of the genetic strategy for targeting excitatory DREADDs to specific neuronal populations in Cre mice.

Wild-type mice received bilateral excitatory DREADD virus (AAV2-CamKIIa-hM3D(Gq)-mCherry) injections into the forelimb region of the MCX. Cervical excitatory DREADD mice received the same excitatory DREADD virus injected bilaterally into the IZ of C5 and C7. Cre mice received excitatory DREADD virus (AAV2-hSyn-hM3D(Gq)-mCherry) injections to activate specific neuronal populations in either the brain stem or spinal cord (Fig 1D). All mice receiving DREADD virus injections received 10 daily CNO injections (i.p.) to enhance neural activity, beginning at least 14 days post-AAV injection to ensure adequate viral transduction.

### Excitatory DREADDs can activate cortico-brain stem-spinal pathways to compensate for CST activity loss and prevent transneuronal degeneration of IN classes

We utilized the biPTX lesion and examined ChAT and Chx10 IN degeneration to determine whether complete loss of CST activity induces transneuronal degeneration (Fig 2A). To confirm that the CST lesions were complete, we determined that PKC gamma staining was absent in the ventral dorsal columns (Jiang et al., 2018). We determined the density of ChAT and Chx10 INs by constructing IN density heatmaps from whole section montages from sections spanning C6-C8 in the spinal cord. ChAT and Chx10 IN counts were determined by IN counts within the region of interest (ROI) in the medial IZ of WT naïve and WT biPTX animals. The ROI is within the field normally receiving CST axon projections (Asante and Martin, 2013), and is expected to show terminal degeneration after biPTX. In WT biPTX mice, there was a decrease in ChAT and Chx10 IN density compared to WT naïve mice (Fig 2B). Compared to WT naive animals, ChAT and Chx10 IN numbers per mm^2^ significantly decreased in WT biPTX animals (Fig 2C, ChAT p<0.0001; Chx10 p=0.0003). These findings extend earlier results in rats with unilateral CST lesions for ChAT INs (Jiang et al., 2018) and show that the major glutaminergic premotor IN class is also subject to degeneration after CST loss.

**Figure 2:**
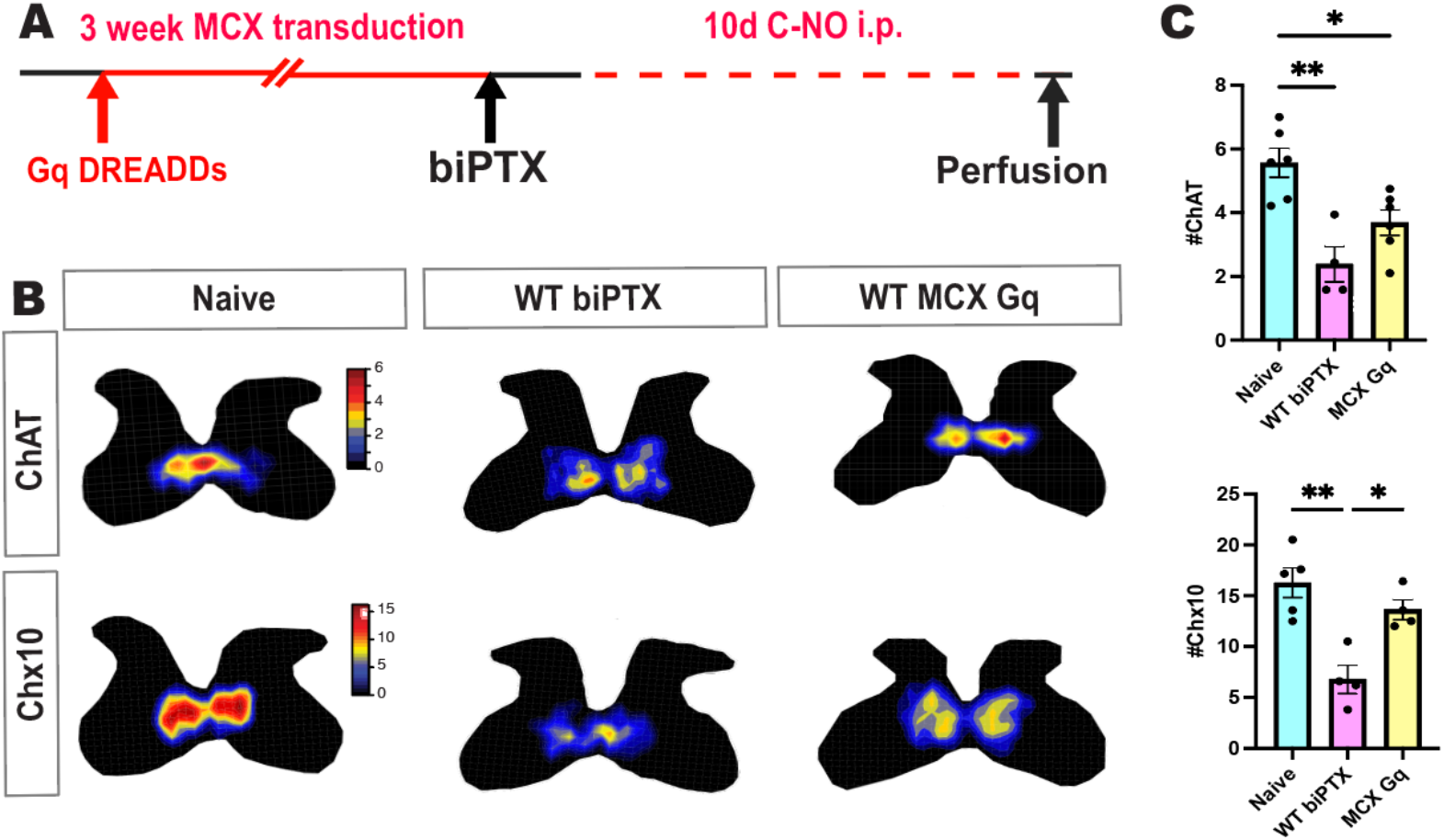
Motor Cortex Excitatory DREADDs. (A) Schematic representation of the experimental timeline showing a 3-week period for motor cortex (MCX) transduction followed by 10 days of clozapine N-oxide (CNO) injections. (B) Heatmaps display the density distribution of ChAT and Chx10 interneurons (INs) in the intermediate zone (IZ) of wild-type (WT) naïve, WT biPTX, and WT MCX Gq mice. (C) Quantification of ChAT and Chx10 INs within the region of interest (ROI) reveals a significant reduction in IN density in WT biPTX mice compared to WT naïve mice (ChAT: p<0.0001; Chx10: p=0.0003). Excitatory DREADD activation in the MCX of WT MCX Gq mice resulted in increased ChAT and Chx10 IN densities compared to WT biPTX mice, with a significant 52% increase in Chx10 IN density (p=0.0172) but a non-significant 44% increase in ChAT IN density (p=0.2357). Error bars represent SEM; each data point represents biological replicates with 10 technical replicates per sample.

We used excitatory DREADDs to activate the motor cortex bilaterally in WT MCX Gq mice after biPTX to determine if enhancing MCX activity can ameliorate IN degeneration without the CST. We hypothesized that MCX excitatory DREADD activation would act through indirect cortico-brain stem-spinal pathways to prevent transneuronal IN degeneration. In WT MCX Gq animals, spinal heatmaps revealed that there was an increase in ChAT and Chx10 IN throughout the IZ compared to that of WT biPTX mice (Fig 2B). Compared to the injury group, Chx10 IN density significantly increased by 52% with WT MCX activation; however, the 44% increase in ChAT IN numbers was not significant (Fig 2C, ChAT p=0.2357; Chx10 p=0.0172). These results show that MCX DREADD activation is capable of ameliorating transneuronal degeneration after complete CST loss.

### Medullary reticular formation (MRF) activation rescues IN classes

A major target of corticofugal projections to the brain stem is the medullary reticular formation (MRF), a major origin of a reticulospinal projection to the cervical cord and an indirect cortico-spinal motor path(Arber, 2012; Esposito et al., 2014). Next, we determined if activation of excitatory VGlut2+ medullary neurons is sufficient to provide neural activity to rescue cervical INs after biPTX. We chose VGlut2 to ensure neurotransmitter specificity of the reticulospinal projection (Esposito et al., 2014). In VGlut2-Cre mice, we targeted bilateral injection of excitatory DREADD virus injections to the ventrolateral medullary reticular formation, including the medullary reticular nucleus, ventral part (MdV), which contains VGlut2+ neurons that are involved with forelimb control (Esposito et al., 2014)(Fig 3A). We visualized DREADD expression (mCherry labeling) at the level of the medullary injection sites to confirm viral transduction (Fig 3B-D). From heatmap topography, we observed an increase in ChAT and Chx10 IN (Fig 3E) in the VGlut2-Cre animals (MRF Gq) compared to the WT biPTX group. To quantify this increase, we found that ChAT and Chx10 IN numbers within the ROI were significantly increased from the WT biPTX group (Fig 2.3F, ChAT p=0.0203; Chx10 p=0.0223).

**Figure 3:**
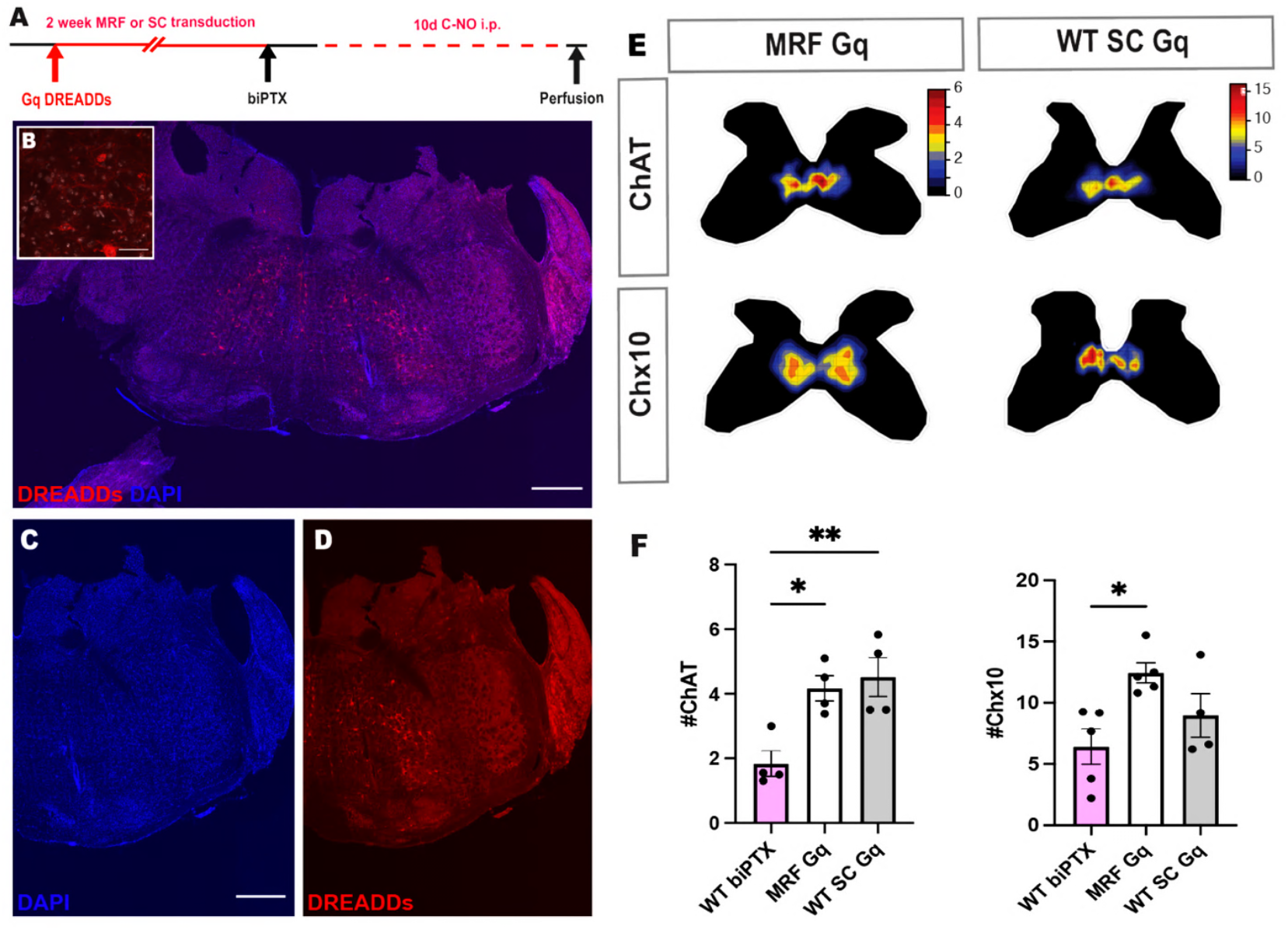
Excitatory DREADDs Bypass the Corticospinal System. (A) Schematic representation illustrating the timeline for bilateral injections of excitatory DREADD virus into the medullary reticular formation (MRF) or spinal cord (SC) in mice. (B) Immunohistochemical analysis showing DREADD (mCherry) expression in the MRF, confirming viral transduction at the injection sites. The inset highlights the high magnification of DREADD-expressing neurons. (C-D) DAPI staining and mCherry labeling of DREADD-expressing neurons in the MRF. (E) Heatmaps showing the density distribution of ChAT and Chx10 interneurons (INs) in the intermediate zone (IZ) of VGlut2-Cre MRF Gq and WT SC Gq mice. (F) Quantification of ChAT and Chx10 INs within the region of interest (ROI) reveals a significant increase in IN density in MRF Gq and WT SC Gq mice compared to WT biPTX mice (ChAT: p=0.0203; Chx10: p=0.0223). Error bars represent SEM; each data point represents biological replicates with 10 technical replicates per sample.

### Generalized neuronal excitation of local spinal circuits ameliorates transneuronal degeneration

We showed that supraspinal neuronal DREADD activation, MCX or MRF, rescues spinal premotor interneurons. We next hypothesized that targeted activation of local spinal circuits can prevent transneuronal degeneration of INs. To test this hypothesis, we used DREADD virus (AAV2-CamKIIa-HM3D(Gq)-mCherry) in the spinal cord of WT mice (WT SC Gq) to activate the local motor circuit after biPTX. Our heatmap results demonstrate that CNO treatment in WT SC Gq animals increased the density of ChAT and Chx10 INs compared with untreated biPTX (see Fig 2B). Compared to the WT biPTX group, Chx10 IN numbers did not return to WT naïve IN numbers after biPTX (Fig 3F, p=0.6531) despite a 41% increase. However, DREADD activation was more effective in ChAT INs, where IN density (numbers of neurons per mm^2^) was not significantly different from the WT SC group (Fig 3C, p=0.0090) and increased by 72%.

This shows that selective activation of spinal cord neurons is effective in supporting IN survival after complete CST loss—with a return to WT baseline for ChAT INs but an incomplete improvement for Chx10 INs. These findings show that different sources of activity can support IN survival after injury, with the caveat that not all sources are necessarily equivalent in efficacy.

### Excitatory DREADD actions abrogate the spinal immune response

To investigate the changes in inflammation associated with CST loss and activity manipulations, we examined microglia activation and phagocytosis through colocalization of Iba-1, a general microglia marker, and CD68, a lysosomal protein marker. We studied this at the 10-day post-injury time point. In the WT naïve animals, microglia tile and extend their processes to survey the environment, indicating a resting state (Fig 4A). In contrast, WT biPTX animals showed a change in microglia morphology, with retracted and thickened processes, indicating an activated state (Fig 4A)(Kozlowski and Weimer, 2012). There was also a significant increase in microglia density (Iba-1 p=0.0006) and phagocytic microglia (Iba-1+CD68+ p<0.0001), indicating microglia proliferation and phagocytosis (Fig 4B).

**Figure 4:**
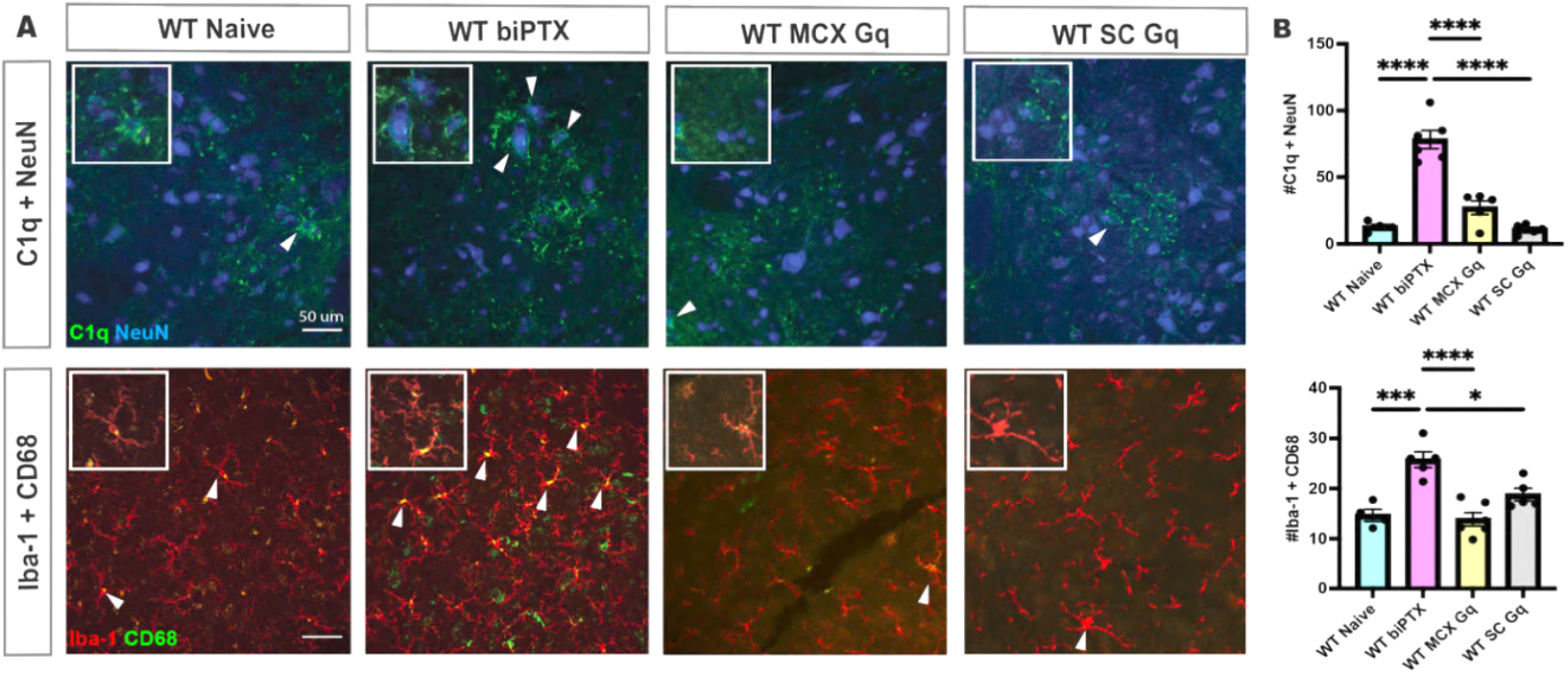
Excitatory DREADDs and Innate Immune Responses. (A) Representative images depict the colocalization of complement protein C1q with the neuronal marker NeuN and microglia marker Iba-1 with lysosomal protein marker CD68. Insets highlight detailed views of the colocalization. Top row images show C1q and NeuN colocalization, indicating complement protein deposits on neurons in WT naïve, WT biPTX, WT MCX Gq, and WT SC Gq mice. Bottom row images show Iba-1 and CD68 colocalization illustrating microglia morphology changes and phagocytic activity in response to CST injury and subsequent activity manipulation. Scale bars = 50 μm. (B) Quantification of C1q+NeuN+ neurons and Iba-1+CD68+ phagocytic microglia numbers. WT biPTX mice show significant increases in both C1q+ neurons and phagocytic microglia compared to WT naïve mice (C1q+NeuN+: p<0.0001; Iba-1+CD68+: p<0.0001). In contrast, WT MCX Gq and WT SC Gq mice, following bilateral excitatory DREADD activation, exhibit significantly reduced numbers of C1q+ neurons and phagocytic microglia, comparable to WT naïve levels (C1q+NeuN+: WT MCX Gq p<0.0001, WT SC Gq p<0.0001; Iba-1+CD68+: WT MCX Gq p<0.0001, WT SC Gq p<0.0001). Error bars represent SEM with each data point representing biological replicates with 10 technical replicates per sample.

These findings led us to examine how innate immune responses may be attuned to changes in neuronal activity after CST injury. Intriguingly, the WT MCX Gq and WT SC Gq groups both show that microglia revert to tiling and show a resting state (Fig 4A). Bilateral excitatory DREADD activation of the motor cortex (WT MCX Gq) or spinal cord (WT SC Gq) after biPTX in wild-type animals significantly decreased both microglia density (Fig 4B; WT MCX Gq p=0.0001 ; WT SC Gq p=0.0123) and phagocytic microglia (WT MCX Gq p<0.0001; WT SC Gq p<0.0001). WT SC Gq and WT MCX Gq groups were not significantly different from WT naïve animals.

Complement protein C1q induces microglia to phagocytose non-apoptotic INs and marks neurons for transneuronal degeneration(Brown and Neher, 2012; Jiang et al., 2018). Our microglia findings prompted us to examine C1q protein changes in WT naïve, WT biPTX, WT MCX Gq, and WT SC Gq models. In WT naïve animals, few to none of the neurons were double-labeled with C1q (C1q+NeuN+) (Fig 4A), consistent with low basal expression in the adult brain (Stephan et al., 2013). Similar to our microglia findings, there was a significant increase in C1q+NeuN+ cells (p<0.0001) in the WT biPTX group when compared to that of the WT naïve group (Fig 4B). Interestingly, in WT MCX Gq or WT SC Gq groups, CNO activation significantly decreased the number of C1q+NeuN+ cells within the ROI (Fig 4C, D, WT MCX Gq p<0.0001; WT SC Gq p<0.0001). Furthermore, the C1q+NeuN+ cell counts in WT MCX Gq and WT SC Gq groups were not significantly different from the WT naïve group (Fig 2.4B, D, WT MCX Gq p>0.9999; WT SC Gq p>0.9999). These findings demonstrate that neural activation ameliorated the innate immune response. C1q+NeuN+ cells in WT MCX Gq and WT SC Gq models returning to WT naïve baseline levels are likely associated with neural activity abrogating C1q production, which mitigates microglia phagocytosis of INs.

### Selective activation of individual spinal IN subclasses rescues the segmental motor circuit

Our finding that spinal DREADD activation abrogates IN C1q expression after CST loss suggests that increased IN activity ameliorates degeneration vulnerability by dampening the immune response. We took advantage of Pitx2 and Chx10 Cre lines for two experiments to elucidate mechanisms further, to determine: 1) if selective activation of one IN class protected both that IN class and the other class, and 2) if most INs that survive by 10 days post-injury do not express C1q, the trigger for non-apoptotic phagocytosis.

To determine if selective activation of one IN class (either Pitx2 or Chx10) protected both that class and the other class, we used Cre-dependent DREADDs (AAV2-hSyn-hM3D(Gq)-mCherry) in either Pitx2-Cre (Pitx2-Cre SC Gq) or Chx10-Cre (Chx10-Cre SC Gq) mice to express DREADDs selectively in Cre-expressing cervical INs (see Methods for injection locations). In this way, we determine if DREADD-Pitx2 IN expression protects ChAT INs only or, additionally, Chx10 INs. Similarly, we determine if DREADD-Chx10 IN expression protects Chx10 INs only or, additionally, ChAT INs. We examined ChAT and Chx10 INs (identified with immunohistochemistry, as in the other analyses) within the ROI.

Compared to WT SC Gq, ChAT and Chx10 IN density was not significantly different in Pitx2-Cre SC Gq and Chx10-Cre SC Gq animals (Fig 5B, Pitx2-Cre SC Gq ChAT p>0.9999, Chx10 p=0.6897; Chx10-Cre SC Gq ChAT p>0.9999, Chx10 p= 0.5785). This suggests that activating one or the other class is as effective as activating all classes.

**Figure 5.**
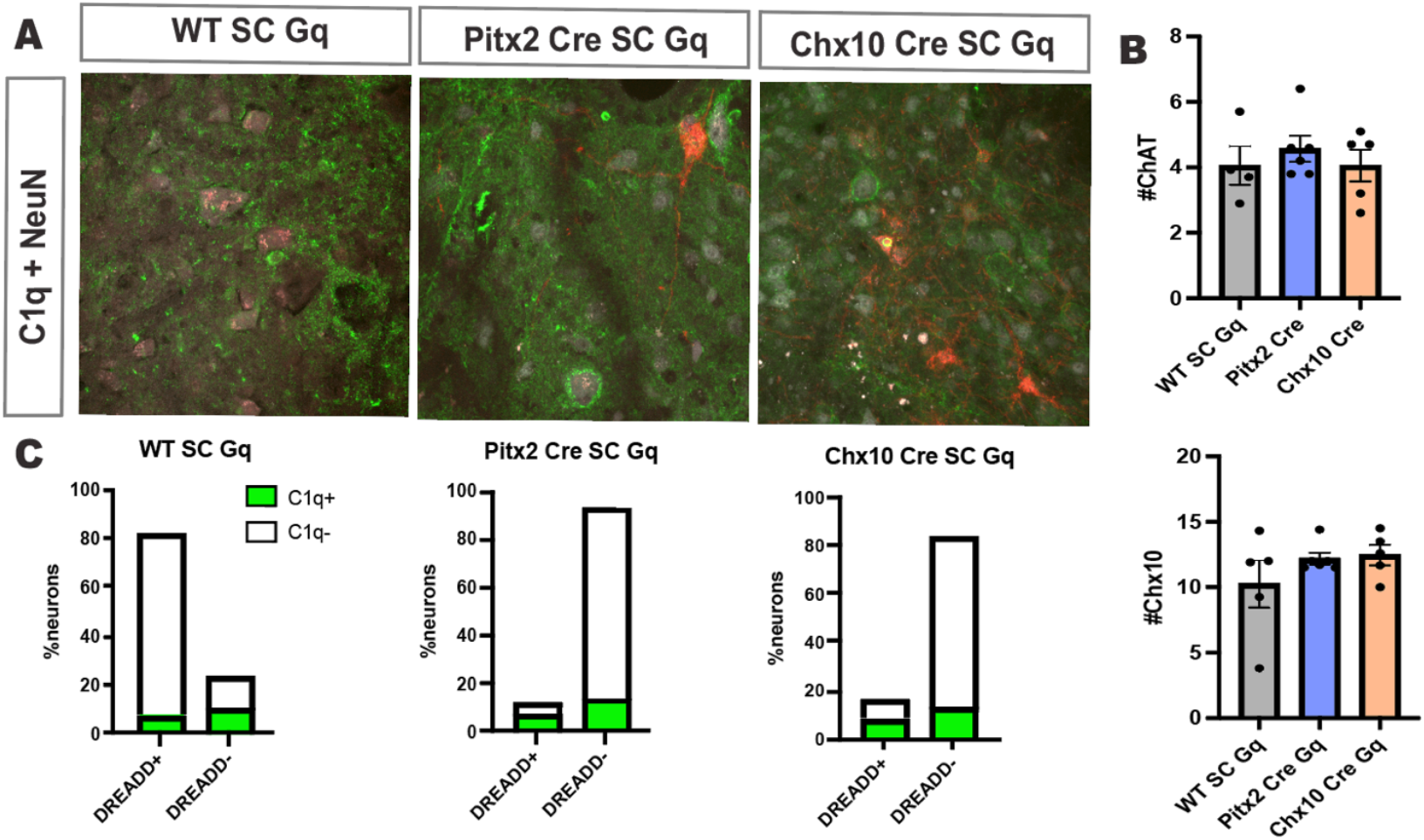
Cell Autonomous Activation on the Immune System. (A) Representative images showing colocalization of complement protein C1q with neuronal marker NeuN in spinal cord sections from WT SC Gq, Pitx2-Cre SC Gq, and Chx10-Cre SC Gq mice. (B) Quantification of ChAT and Chx10 interneurons (INs) within the region of interest (ROI) across different experimental groups shows no significant differences. Error bars represent SEM. (C) Percentage of C1q+ and C1q-neurons among DREADD+ and DREADD-neurons in WT SC Gq, Pitx2-Cre SC Gq, and Chx10-Cre SC Gq mice, indicating that most DREADD-neurons are C1q-.

Compared with generalized SC IN DREADD activation (WT SC Gq), selective DREADD IN activation resulted in no difference in total IN numbers. Pitx2-Cre SC Gq and Chx10-Cre SC Gq animals had significantly increased ChAT interneuron numbers—44% and 41%, respectively—displaying activity-dependent IN rescue. These findings support our hypothesis that activating individual IN classes indirectly activates the local spinal circuit to ameliorate transneuronal degeneration more generally.

We next determined if DREADD-activated INs that survive by 10 days post-injury do not express C1q, suggesting that they are less vulnerable to degeneration. We counted the number of DREADD+ and DREADD-neurons with and without C1q expression (2.5C). For this analysis, we did not distinguish IN subclasses. For generalized SC IN DREADD activation (Fig 5C, WT SC Gq), among the DREADD+ neurons (77%), the largest class was C1q-(74%). The DREADD-INs (24%) showed a slight preponderance for C1q-, suggesting that DREADD-activated neurons can indirectly activate DREADD-neurons to quell the inflammatory response (Fig 5C).

For both the Pitx2 and Chx10 classes, the number of DREADD+ neurons was small by comparison to DREADD-(i.e., non-Cre vs Cre ; Pitx2: 16% versus 84%; Chx10: 15% versus 85%). Not surprisingly, the largest group comprised DREADD-, and most of these neurons were C1q-, suggesting that the small number of DREADD+ Pitx2 and Chx10 INs were capable of activating the general population. Indeed, as Fig 5B shows, selective activation of either subclass was as effective in rescuing spinal INs as generalized DREADD activation. These findings suggest that a small number of activated spinal INs can protect the general population through divergent excitatory interactions to ameliorate the inflammatory response.

### Preventing transneuronal degeneration is associated with functional recovery

Our finding that activating spinal INs excites the general spinal circuit for divergent excitatory interactions prompted us to investigate whether modulating neural activity at the cellular level to prevent transneuronal degeneration is associated with function recovery. To answer this question, we used the grip strength task. We chose this task to study functional recovery since the CST contributes to dexterous digit movements(Anderson et al., 2004), and the task is a voluntary reach-to-grasp response. Grip strength (peak tension force) was assessed before biPTX (day 0) and 1-day post-injury (dpi), 3dpi, 5dpi, and 10dpi. Grip force in WT naive mice is 4.20 g, significantly reduced after biPTX (Fig 6A-B, p <0.0001). We determined if DREADD activation of the MRF, Pitx2, and Chx10 improved grip strength after biPTX.

**Figure 6:**
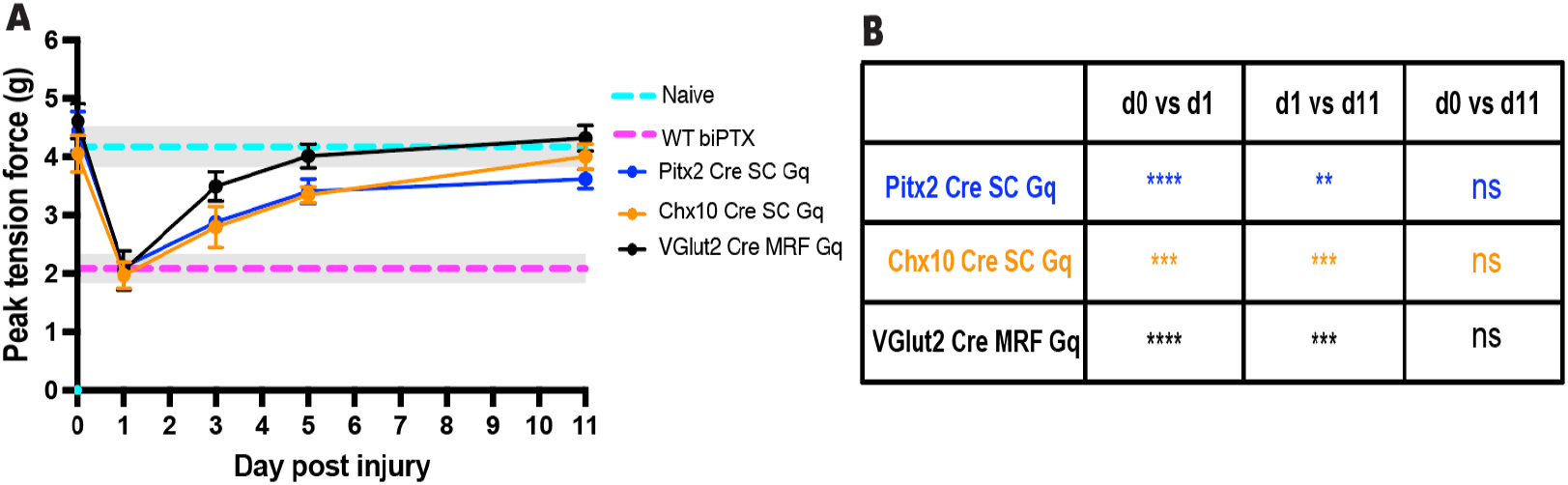
Grip strength recovery following CST elimination and modulation of neural activity. (A) Peak tension force (grip strength) was assessed in WT naïve, WT biPTX, Pitx2-Cre SC Gq, Chx10-Cre SC Gq, and VGlut2-Cre MRF Gq mice before injury (day 0) and at various time points post-injury (1dpi, 3dpi, 5dpi, and 11dpi). Systemic administration of CNO was used to modulate neural activity via DREADDs, with grip strength measured during the washout period. (B) Statistical comparison of grip strength across different time points. Error bars represent SEM.

The non-DREADD + CNO group consistently replicated the baseline grip strength effects in the DREADD + CNO transfected animals, and statistical analysis revealed no significant differences between the groups (data not shown, p>0.999). Grip strength was assessed 1day post-injury (dpi), 3dpi, 5dpi, and 10dpi before CNO injections to ensure readouts from the assay resulted from a net increase in neural activity and not the actions of CNO.

VGlut2-Cre MRF Gq, Pitx2-Cre SC Gq, and Chx10-Cre SC Gq animals all demonstrated a significant decrease in grip strength peak tension force on 1dpi (Fig 6A-B; VGlut2-Cre MRF Gq p<0.0001; Pitx2-Cre SC Gq p<0.0001; and Chx10-Cre SC Gq p=0.0003). We observed a gradual and progressive increase in grip strength across all Cre models after biPTX until the experiment’s endpoint. Grip strength significantly improved between 1dpi and 11dpi (Fig 6A-B, VGlut2-Cre MRF Gq p=0.0002, Pitx2-Cre SC Gq p=0.0012, and Chx10-Cre SC Gq p=0.0003) and there were no significant differences in peak tension force on day 0 and by 11dpi (Fig 6A-B, VGlut2-Cre MRF Gq p>0.9999, Pitx2-Cre SC Gq p=0.0931, and Chx10-Cre SC Gq p>0.9999). These results show that modulating neural activity improves functional recovery after complete CST elimination.

## Discussion

This study explores the critical role of neural activity in maintaining spinal premotor circuits and motor function following CST injury. Spinal cord INs integrate proprioceptive and descending control signals, ultimately driving motor neuron activation (Baldissera et al., 1981; Lemon, 2008). The vulnerability of these INs to transneuronal degeneration in the absence of CST activity, either due to complete lesion or loss of MCX activity using inhibitory DREADDs, underscores the contribution of an activity dependency to maintaining their viability. Notably, spinal INs can be protected from degeneration through activity-based interventions at multiple levels of the neuraxis. Without any CST connections in the spinal cord after bilateral PTX, excitatory DREADD activation of the MCX or spinal cord prevented transneuronal degeneration, depending on the activation site. Further, using mouse Cre lines to target ChAT (Zagoraiou et al., 2009)and Chx10 (Ueno et al., 2018)interneurons—as well as glutamatergic reticular formation neurons, including the reticulospinal tract (Esposito et al., 2014)— transneuronal degeneration was abrogated, and this prevented the loss of forepaw grip strength after bilateral PTX. Our findings highlight the potential for therapeutic strategies to maintain IN populations and restore spinal circuits’ function following CST injury.

### Transneuronal degeneration results from the loss of neural activity, not CST trophic signals

While we initially hypothesized that transneuronal degeneration is primarily driven by the loss of trophic interactions from CST neurons, based on our earlier studies showing failure of ChAT interneuron development with postnatal motor cortex inactivation(Chakrabarty et al., 2009; Friel et al., 2012), our results demonstrate that transneuronal degeneration results from the loss of segmental neural activity rather than CST-specific trophic signals. A key observation is the significant reduction in transneuronal IN degeneration when selectively activating the forelimb MCX despite the complete loss of CST connections. This showed that the motor cortex acted through spared connections, especially those of the brain stem pathways to the spinal cord. This result prompted us to investigate downstream sources of neural activity that can maintain IN viability.

### Dissection of subcortical circuit contributions reveals the generalized role of activity in transneuronal degeneration

Our study aimed to identify the subcortical circuits involved in mediating motor cortex activation after CST lesion, emphasizing the role of activity at the segmental level in transneuronal degeneration. Using DREADDs for their precision and control in manipulating neural circuits by visualizing the neurons that were either inactivated or activated, we aimed to understand better the effects of activity modulation on INs after CST loss. We hypothesized that indirect MCX to spinal cord pathways could substitute neural activity signals to spinal INs, thereby preventing transneuronal degeneration. Our study focused on the reticulospinal pathway due to its evolutionary conservation across mammals by targeting the ventrolateral medullary reticular formation (MRF), where the MdV is located (Matsuyama et al., 2004; Liang et al., 2016; Asboth et al., 2018; Engmann et al., 2020; Usseglio et al., 2020; Olivares-Moreno et al., 2021). While our findings indicate that MRF activation significantly ameliorated transneuronal degeneration of specific IN populations, complementary methods, such as dual viral constructs to target reticulospinal neurons for DREADD activation selectively, would enhance our understanding further.

### The dynamic interplay between neuronal activity and the immune responses for the regulation of neuroinflammation

Microglia, the resident immune cells of the CNS, play a role in neuroinflammation regulation (Jebelli et al., 2015; Szepesi et al., 2018; Borst et al., 2021). We propose that normal resting-state activity maintains microglial surveillance, whereas the activity loss triggers swift microglial activation. Over an extended post-lesion timeframe, we show an inverse correlation between the microglial activation state and the level of neuronal activity. Specifically, spinal microglial activation increases with inhibitory DREADD action in the motor cortex in intact mice, transitioning to a surveillance state upon restoration through excitatory DREADDs, as shown after biPTX. This demonstrates the bidirectional relationship between neuronal activity and immune responses in the CNS.

Excitatory DREADD activation of motor cortex or reticular formation neurons achieved transition to the spinal microglial surveillance state. This underscores the influence of conducted activity—including indirect corticofugal pathway activation or direct activation of RF neurons— to compensate for the baseline activity of the CST prior to injury. Crucially, reducing neuronal C1q production, driven by increased spinal cord activity from different sources, is an effective strategy for preserving IN survival by mitigating the trigger for microglial phagocytosis and may be replicated by spinal neuromodulation (Jiang et al., 2018). In transneuronal degeneration, our results align with findings in rats, where CST loss initially elevated C1q and subsequently abrogated as vulnerable neurons degenerated (Jiang et al., 2018). To address how neuromodulation drives microglial activation independent of its direct actions on neurons to increase activity, our data support that spinal DREADD activation replicates actions of ts-DCS neuromodulation. These data provide a potential mechanism that explains the observed.

**Table 1:**
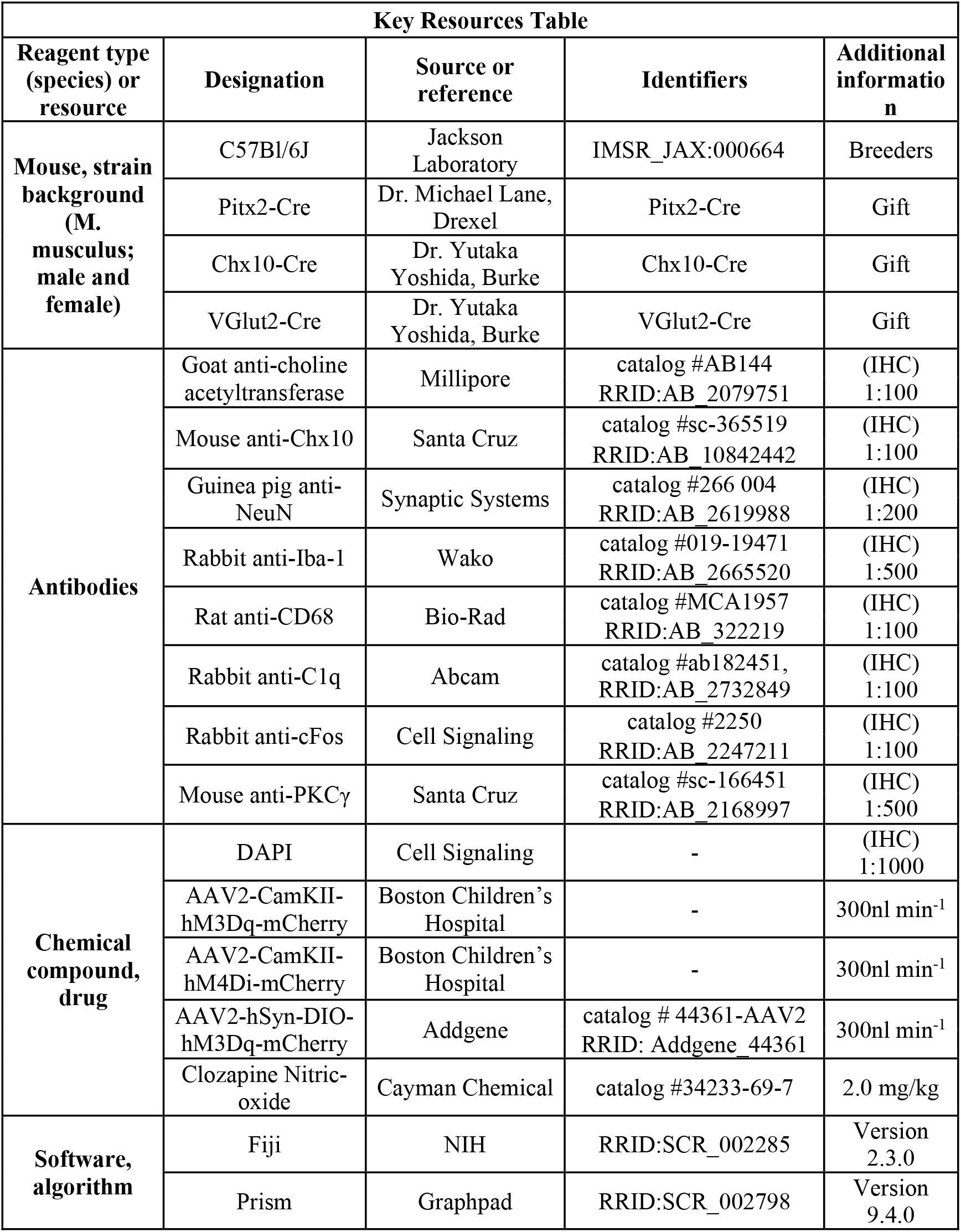
Key Resources Table.

**Table 2:**
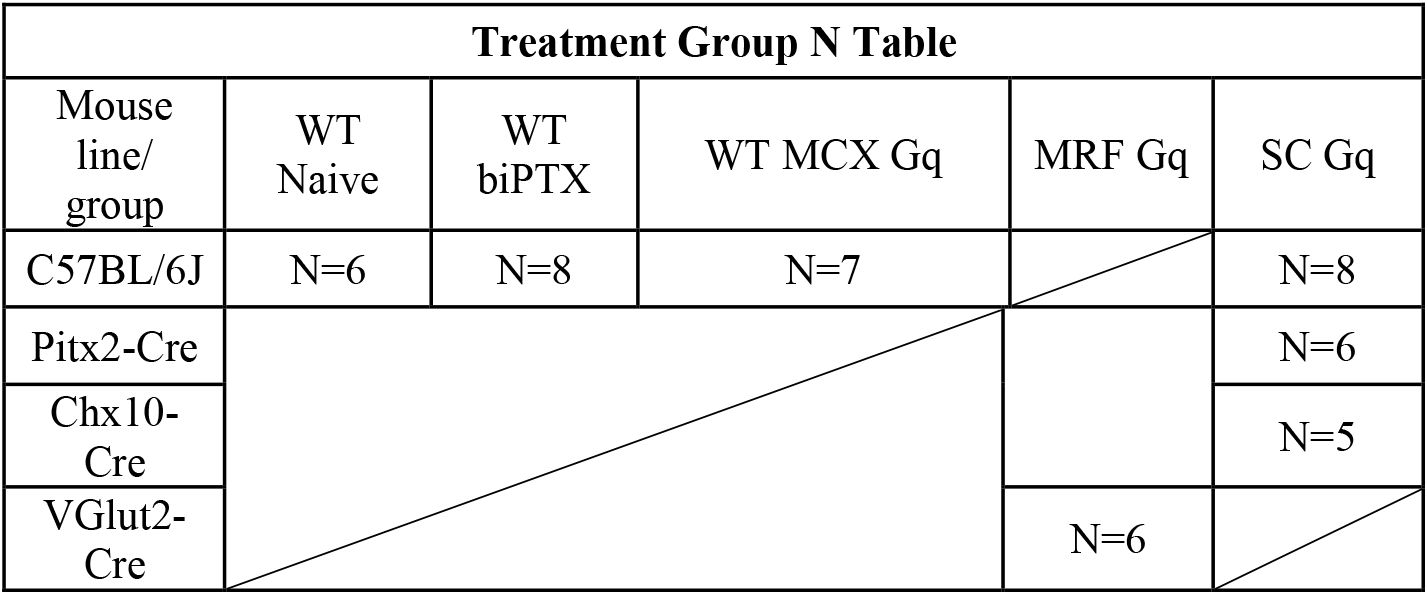
Treatment Group N Table.

## Acknowledgements

We thank Dr Michael Lane, Drexel University School of Medicine, and Dr Gareth Miles, University of St. Andrews, for the generous gift of Pitx2-Cre mice and XiuLi Wu for histology.

The authors have no competing interests to declare.

## References

Aartsma-Rus A, van Putten M (2014) Assessing functional performance in the mdx mouse model. J Vis Exp.

Amer A, Martin JH (2022) Repeated motor cortex theta-burst stimulation produces persistent strengthening of corticospinal motor output and durable spinal cord structural changes in the rat. Brain Stimul 15:1013–1022.

Anderson KD, Abdul M, Steward O (2004) Quantitative assessment of deficits and recovery of forelimb motor function after cervical spinal cord injury in mice. Exp Neurol 190:184–191.

Arber S (2012) Motor circuits in action: specification, connectivity, and function. Neuron 74:975–989.

Asante CO, Martin JH (2013) Differential joint-specific corticospinal tract projections within the cervical enlargement. PLoS One 8:e74454.

Asboth L, Friedli L, Beauparlant J, Martinez-Gonzalez C, Anil S, Rey E, Baud L, Pidpruzhnykova G, Anderson MA, Shkorbatova P, Batti L, Pages S, Kreider J, Schneider BL, Barraud Q, Courtine G (2018) Cortico-reticulo-spinal circuit reorganization enables functional recovery after severe spinal cord contusion. Nat Neurosci 21:576–588.

Baldissera F, Hultborn H, Illert M (1981) Integration in spinal neuronal systems. In: Handbook of Physiology, Section I: The Nervous System, vol II, Motor Control (Brooks VB, ed), pp 509–596. Bethesda: American Physiological Society.

Bareyre FM, Kerschensteiner M, Misgeld T, Sanes JR (2005) Transgenic labeling of the corticospinal tract for monitoring axonal responses to spinal cord injury. Nat Med 11:1355–1360.

Borst K, Dumas AA, Prinz M (2021) Microglia: Immune and non-immune functions. Immunity 54:2194–2208.

Brosamle C, Schwab ME (2000) Ipsilateral, ventral corticospinal tract of the adult rat: ultrastructure, myelination and synaptic connections. J Neurocytol 29:499–507.

Brown GC, Neher JJ (2012) Eaten alive! Cell death by primary phagocytosis: ‘phagoptosis’. Trends Biochem Sci 37:325–332.

Brown GC, Neher JJ (2014) Microglial phagocytosis of live neurons. Nat Rev Neurosci 15:209–216.

Brown GC, Vilalta A, Fricker M (2015) Phagoptosis - Cell Death By Phagocytosis - Plays Central Roles in Physiology, Host Defense and Pathology. Curr Mol Med 15:842–851.

Chakrabarty S, Shulman B, Martin JH (2009) Activity-dependent codevelopment of the corticospinal system and target interneurons in the cervical spinal cord. J Neurosci 29:8816–8827.

Dougherty KJ, Kiehn O (2010) Firing and cellular properties of V2a interneurons in the rodent spinal cord. J Neurosci 30:24–37.

Engmann AK, Bizzozzero F, Schneider MP, Pfyffer D, Imobersteg S, Schneider R, Hofer AS, Wieckhorst M, Schwab ME (2020) The Gigantocellular Reticular Nucleus Plays a Significant Role in Locomotor Recovery after Incomplete Spinal Cord Injury. J Neurosci 40:8292–8305.

Esposito MS, Capelli P, Arber S (2014) Brainstem nucleus MdV mediates skilled forelimb motor tasks. Nature 508:351–356.

Friel K, Chakrabarty S, Kuo HC, Martin J (2012) Using Motor Behavior during an Early Critical Period to Restore Skilled Limb Movement after Damage to the Corticospinal System during Development. J Neurosci 32:9265–9276.

Hagglund M, Dougherty KJ, Borgius L, Itohara S, Iwasato T, Kiehn O (2013) Optogenetic dissection reveals multiple rhythmogenic modules underlying locomotion. Proc Natl Acad Sci U S A 110:11589–11594.

Jebelli J, Su W, Hopkins S, Pocock J, Garden GA (2015) Glia: guardians, gluttons, or guides for the maintenance of neuronal connectivity? Ann N Y Acad Sci 1351:1–10.

Jiang Y-Q, Sarkar A, Amer A, Martin JH (2018) Transneuronal Downregulation of the Premotor Cholinergic System After Corticospinal Tract Loss. Journal of Neuroscience 38:83298344.

Kozlowski C, Weimer RM (2012) An automated method to quantify microglia morphology and application to monitor activation state longitudinally in vivo. PLoS One 7:e31814.

Lemon RN (2008) Descending pathways in motor control. Annu Rev Neurosci 31:195–218.

Liang H, Watson C, Paxinos G (2016) Terminations of reticulospinal fibers originating from the gigantocellular reticular formation in the mouse spinal cord. Brain Struct Funct 221:1623–1633.

Matsuyama K, Mori F, Nakajima K, Drew T, Aoki M, Mori S (2004) Locomotor role of the corticoreticular-reticulospinal-spinal interneuronal system. Prog Brain Res 143:239–249.

Olivares-Moreno R, Rodriguez-Moreno P, Lopez-Virgen V, Macias M, Altamira-Camacho M, Rojas-Piloni G (2021) Corticospinal vs Rubrospinal Revisited: An Evolutionary Perspective for Sensorimotor Integration. Frontiers in neuroscience 15:686481.

Roth BL (2016) DREADDs for Neuroscientists. Neuron 89:683–694.

Serradj N, Paixao S, Sobocki T, Feinberg M, Klein R, Kullander K, Martin JH (2014) EphA4-mediated ipsilateral corticospinal tract misprojections are necessary for bilateral voluntary movements but not bilateral stereotypic locomotion. J Neurosci 34:5211–5221.

Serradj N, Marino F, Moreno-Lopez Y, Bernstein A, Agger S, Soliman M, Sloan A, Hollis E (2023) Task-specific modulation of corticospinal neuron activity during motor learning in mice. Nat Commun 14:2708.

Stephan AH, Madison DV, Mateos JM, Fraser DA, Lovelett EA, Coutellier L, Kim L, Tsai HH, Huang EJ, Rowitch DH, Berns DS, Tenner AJ, Shamloo M, Barres BA (2013) A dramatic increase of C1q protein in the CNS during normal aging. J Neurosci 33:13460–13474.

Szepesi Z, Manouchehrian O, Bachiller S, Deierborg T (2018) Bidirectional Microglia-Neuron Communication in Health and Disease. Front Cell Neurosci 12:323.

Ueno M, Nakamura Y, Li J, Gu Z, Niehaus J, Maezawa M, Crone SA, Goulding M, Baccei ML, Yoshida Y (2018) Corticospinal Circuits from the Sensory and Motor Cortices Differentially Regulate Skilled Movements through Distinct Spinal Interneurons. Cell Rep 23:1286–1300 e1287.

Usseglio G, Gatier E, Heuze A, Herent C, Bouvier J (2020) Control of Orienting Movements and Locomotion by Projection-Defined Subsets of Brainstem V2a Neurons. Curr Biol 30:4665–4681 e4666.

Yang L, Martin JH (2023) Effects of motor cortex neuromodulation on the specificity of corticospinal tract spinal axon outgrowth and targeting in rats. Brain Stimul 16:759–771.

Zagoraiou L, Akay T, Martin JF, Brownstone RM, Jessell TM, Miles GB (2009) A cluster of cholinergic premotor interneurons modulates mouse locomotor activity. Neuron 64:645–662.

